# Hop stunt viroid infection alters host heterochromatin

**DOI:** 10.1101/2023.12.12.571286

**Authors:** Joan Marquez-Molins, Jinping Cheng, Julia Corell-Sierra, Vasti Thamara Juarez-Gonzalez, Pascual Villalba-Bermell, Maria Luz Annacondia, Gustavo Gomez, German Martinez

## Abstract

Viroids are pathogenic non-coding RNAs that completely rely on their host molecular machinery to accomplish their life cycle. Several interactions between viroids and their host molecular machinery have been identified, including an interference with epigenetic mechanisms such as DNA methylation. Despite this, whether viroids influence changes in other epigenetic marks such as histone modifications remained unknown. Epigenetic regulation is particularly important during pathogenesis processes because it might be a key regulator of the dynamism of the defense response. Here we have analyzed the changes taking place in *Cucumis sativus* facultative and constitutive heterochromatin during hop stunt viroid (HSVd) infection using chromatin immunoprecipitation (ChIP) of the two main heterochromatic marks: H3K9me2 and H3K27me3. We find that HSVd infection is associated with changes in both H3K27me3 and H3K9me2, with a tendency to decrease the levels of repressive epigenetic marks through infection progression. These epigenetic changes are connected to the transcriptional regulation of their expected targets, genes and transposable elements. Indeed, several genes related to the defense response are targets of both epigenetic marks. Our results highlight another host regulatory mechanism affected by viroid infection, providing further information about the complexity of the multiple layers of interactions between pathogens/viroids and hosts/plants.

## INTRODUCTION

Viroids are fascinating biological entities characterized by their extremely simple genomes, short (between 200 and 400-nt) circular non-coding RNAs, that are only pathogenic to plants (1,2). Classified as sub-viral pathogens, viroids can be subdivided into two families according to their ability to replicate in the nucleus (*Pospiviroidae*) or the chloroplast (*Avsunviroidae*) (3,4). Due to their remarkably simple genomic organization, viroids must interact with the molecular machinery of the plant cell to fulfill every aspect of their life cycle. As a consequence of this extremely close interaction with their host, viroids induce developmental defects that are identified as symptomatology and are very similar to symptoms caused by viruses (1,2). In general, symptoms expression is the manifestation of the alteration in the development/defense tradeoff, which, when disturbed, is detrimental to plant growth (5). Due to the importance of maintaining such balance, the defense response is controlled by multiple overlapping mechanisms. Several pieces of evidence indicate that epigenetic pathways are one of the key regulators of stress-associated transcriptional reprogramming (6–8). The dynamic nature of the changes introduced by these mechanisms and their known role in the interpretation of environmental and developmental cues in the cellular program makes them good candidates as master regulators of the response to stress (6–8). Epigenetic mechanisms comprise both DNA methylation and histone modifications, both of which play a role in the orchestration of genome stability and transcriptional programs (9).

DNA methylation is the best understood epigenetic mark. It is mediated by the covalent modification of the residue cytosine (C) of the genomic DNA with the addition of a methyl group (9). DNA methylation actively targets transposable elements (TEs, where it accumulates in three sequence contexts: CG, CHG and CHH, being H any nucleotide other than C), being also able to target genes similarly to TEs (accumulating in three sequence context) or exclusively marking the gene body (accumulating only in the CG context) (9,10). This targeting induces the silencing of both TEs and genes (11). In addition to DNA methylation, plants have another conserved epigenetic mechanism consisting of posttranslational modifications of the histone tails that are part of the nucleosome (12). These modifications define the interactions between neighboring nucleosomes, leading to the formation of two different levels of chromatin compaction: euchromatin and heterochromatin, that contain loosely or highly compacted nucleosomes, the latest being recalcitrant to RNA II polymerase-mediated transcription (13). Heterochromatin can be further divided into two different structural forms: facultative and constitutive, which are dynamic or not to developmental/environmental signals (13,14). Interestingly, both forms of heterochromatin have different genomic targets and deposition machinery.

Facultative heterochromatin is marked by the trimethylation of the lysine 27 of histone 3 (H3K27me3), which is located in gene-rich genomic regions within the arms of the chromosomes (15,16). H3K27me3 is controlled by the protein complexes Polycomb repressive complex 1 and 2 (PRC1 and PRC2) and marks the promoters and transcriptional start sites of specific genes (17–20). Recent evidences indicate that, additionally, H3K27me3 controls the formation of repressive chromatin hubs (21). H3K27me3 is a key regulatory epigenetic mark that controls a myriad of developmental events including flowering time, circadian clock, sensing of temperature and gene imprinting (22–28). On the other hand, constitutive heterochromatin is marked by several histone marks, including H3K27me1, H3K9me1 and H3K9me2 (29). This type of heterochromatin is enriched in centromeric and pericentromeric regions where it targets TEs and other repetitive sequences. H3K9 methylation is mediated by members of the SUVR class (SU(VAR)3–9 homologs (SUVH) and SU(VAR)3–9 related proteins) (29). In plants, H3K9 methylation is mechanistically connected to DNA methylation, via the ability of SUVH members to recognize non-CG methylation (15,16,18), which creates a feedback loop with the methyltransferase CMT3, which can recognize H3K9 methylation (30). Alternatively, H3K9 methylation can also be established without a premethylated DNA state (31).

Both epigenetic mechanisms (DNA methylation and histone marks) have been identified as active players in the response against stress in both genetic (32–34) and genome-wide (35–38) studies. Particularly, DNA methylation is a dynamic stress-responsive epigenetic mark under both biotic and abiotic stresses (39–51) that can even be transmitted transgenerationally (52,53). Additionally, histone marks have been proposed to play a role in regulation of the stress-associated transcriptional reprogramming (35,54–60) although their dynamism and role under stress in plants has not been studied in detail until recently (37,38,61).

Nuclear viroid infection has been reported to induce both the hypomethylation of ribosomal repeats and TEs in several species (46–48,62), and the hypermethylation of host genes (62,63) and co-infective DNA viruses (64). Whether these changes are connected to histone marks is completely unexplored. Interestingly, viroids can interfere with Histone Deacetylase 6 (HDA6) suggesting a potential effect over histone mark homeostasis (65). Despite these evidences, there is a substantial lack of genome-wide studies identifying both the targets and extent of epigenetic changes (especially histone marks) associated to viroid pathogenesis. Here, to understand the dynamism of repressive histone marks during viroid infection, we studied the genome-wide presence of the repressive histone marks H3K9me2 and H3K27me3 and their correlation with both DNA methylation and transcription in response to Hop stunt viroid (HSVd) infection in cucumber, at two different time points. We found that HSVd leads to a re-organization of H3K9me2 and H3K27me3 characterized by decrease of the presence of both marks over repeats and different patterns of dynamism over genes. Importantly, H3K27me3 is globally correlated with transcriptional changes, including several stress-responsive genes that might be important to respond to viroid infection. In summary, our data offers a novel perspective about viroid-host interactions and the interplay between heterochromatin dynamism and the response to adverse environmental conditions in plants.

## MATERIAL AND METHODS

### Plant material and HSVd infection

*Cucumis sativus* (ecotype Marketer) were sown into potting soil (P-Jord, Hasselfors Garden, Örebro, Sweden) into plastic pots (9 × 9 × 7 cm) with one plant per pot at a temperature of 30°C and 60% relative humidity. Plants were grown under a 16 h: 8 h, light : dark photoperiod. The light was provided by FQ, 80 W, Hoconstant lumix (Osram, Munich, Germany) with a light intensity of 220 μmol photons m^−2^ s^−1^. HSVd infection was performed at the two-cotyledon stage. Seedlings were agroinfiltrated with an infectious HSVd clone or an empty vector (mock plants)(66). Samples were collected at 10 and 27 dpi.

### Chromatin Immunoprecipitation (ChIP) sequencing libraries preparation and sequence analysis

First, 500 mg of systemic leaves were chemically cross-linked using 1% formaldehyde. Then, nuclei were isolated from cross-linked material following a standard nuclei isolation protocol based on sucrose gradients as previously described (67). The resuspended nuclei pellets were sonicated for 9 cycles of 20 s On and 45 s Off at 4 °C and high power to obtain the chromatin. Afterward, the Immunoprecipitation (IP) was performed following a standard IP protocol and using the following antibodies: H3 (Reference: 07-690, Merck), H3K9me2 (Reference: pAb-060-050, Diagenode) and H3K27me3 (Reference: 07-449, Merck). The resulting immunocomplexes were purified with the GeneJET PCR Purification Kit (Thermo Fisher), following the manufacturer’s instructions. Finally, DNA libraries were prepared using the NEBNext® Ultra™ II DNA Library Prep Kit for Illumina® (New England Biolabs) and each individual library was barcoded using the NEBNext® Multiplex Oligos for Illumina® kit (New England Biolabs).

ChIP libraries of two bioreplicates per condition were obtained from the immunoprecipitated DNA and sequenced as paired-end 150 bp fragments in an Illumina Novaseq 6000 at Novogene (Beijing, China). The obtained raw reads were trimmed using Trimgalore 0.6.1 to remove the adapter sequences and 10 bases from 5’ ends. For genome-wide distribution analysis sequences were aligned to the *Cucumis sativus* “Chinese long” version 3 genome (68) using bowtie2 with default parameters. BAM files were filtered for unique reads using the parameter -q10 and replicates were merged using samtools (69). Genome coverage was calculated as the log2 fold change of the ratio between the coverage of H3K9me2 or H3K27me3 to the coverage of H3 using deeptools2 (70). For genome-wide profile images, the RPKM-normalized value of H3 was subtracted from the RPKM value of either H3K9me2 or H3K27me3. Values for specific regions and quantitative analysis were retrieved using mapbed from bedtools (71).

For peak identification, reads were aligned to the *Cucumis sativus* “Chinese long” version 3 genome using bowtie2, with the options --no-mixed --no-discordant. BAM files were filtered for unique reads and high-quality alignments using the parameters -q 10 and -F 256 using samtools. Peak calling was performed using Sicer2 each sample to its respective H3 control with the parameters window size 200, fragment size 150, effective genome fraction 0.74, false discovery rate 0.01, false discovery rate of 0.01 and a gap size of 600bp. Peak location and overlap was compared using the intersect tool from bedtools with a minimum overlap of 1bp. Only peaks shared between the two replicates were considered true peaks for that specific treatment. Shared peaks were compared between samples using the intersect tool from bedtools to determine gain and loss peaks.

### Gene ontology (GO) term analysis

GO term analysis was performed using the GO annotation for the *Cucumis sativus* “Chinese long” version 3 (http://cucurbitgenomics.org/organism/20) (72). GO categories were simplified to the GO Slim Classification for Plants.

### DNA methylation and RNA sequencing re-analysis

DNA methylation values were obtained from the previous analysis in Marquez-Molins *et al*. (2023). RNA sequencing libraries from the same study (62) were filtered to infer the expression level of repeats. In brief, the count reads per repeat were obtained using HTSeq-COUNTS with the following parameters: --mode union --stranded no --minequal 0 and --nonunique all. The obtained count tables were used in DESeq2 (73) to infer significant expression with fit type set to parametric. All these tools were used on the Galaxy platform (74). Volcano plots were created using the R package ggplot2 (75).

## RESULTS

### Overview of *Cucumis sativus* heterochromatin

Previous studies have shown that alteration in DNA methylation patterns is dynamic under HSVd infection and an important component of the reprogramming of the defense response against this sub-viral pathogen (46–48,62). The involvement of other epigenetic mechanisms such as histone marks that could respond to viroid infection could be inferred from their direct interaction with HDA6 (65) but their overall genome-wide changes are not understood. Heterochromatin, composed of facultative and constitutive regions, is an important genomic structure controlling the stability of the genome and the expression of genes. To understand the dynamism and genome-wide changes taking place in heterochromatin under viroid infection, we performed ChIP-sequencing for the two best studied heterochromatic histone modifications H3K27me3 and H3K9me2 in cucumber plants infected with HSVd at two different time points: 10 days post-infection (dpi, onset of infection) and 27 dpi (development of symptoms) equivalent to the time points used in Marquez-Molins *et al*. (2023).

Our ChIP offered a unique opportunity to study heterochromatin in cucumber, which was not yet described. In mock tissues, the distribution of the two histone marks showed a similar pattern to the one observed in other plants such as *Arabidopsis thaliana* (*76*). H3K9me2 accumulated to a higher level in centromeric regions enriched in repeats, while H3K27me3 was enriched in the arms of the chromosomes where the density of repeats is lower (Fig 1A). As expected from this pattern, genes are preferentially marked by H3K27me3, while repeats are usually marked by H3K9me2 (Fig 1B). Targets of both marks included well-studied epigenetically regulated examples such as an FLC homolog in cucumber (CsaV3_3G016650) targeted by H3K27me3, and a long Gypsy TE targeted by H3K9me2 (Sup Fig 1). The enrichment of H3K9me2 among repeats was more obvious among TEs, with members of the Gypsy, LINE, and Copia retrotransposon superfamilies showing higher values of this repressive mark compared to other DNA transposon superfamilies such as hAT-AC, MuDR and En-SPM (Fig 1C). TEs that presented high values of H3K9me2 had characteristics of heterochromatic entities, such as longer length and closer distance to the centromere (Sup Fig 2A-B). As expected, longer TEs tend to have higher values of H3K9me2 (Sup Fig 2C), a characteristic that is not present for H3K27me3 and genes (Sup Fig 2D).

**Figure 1.**
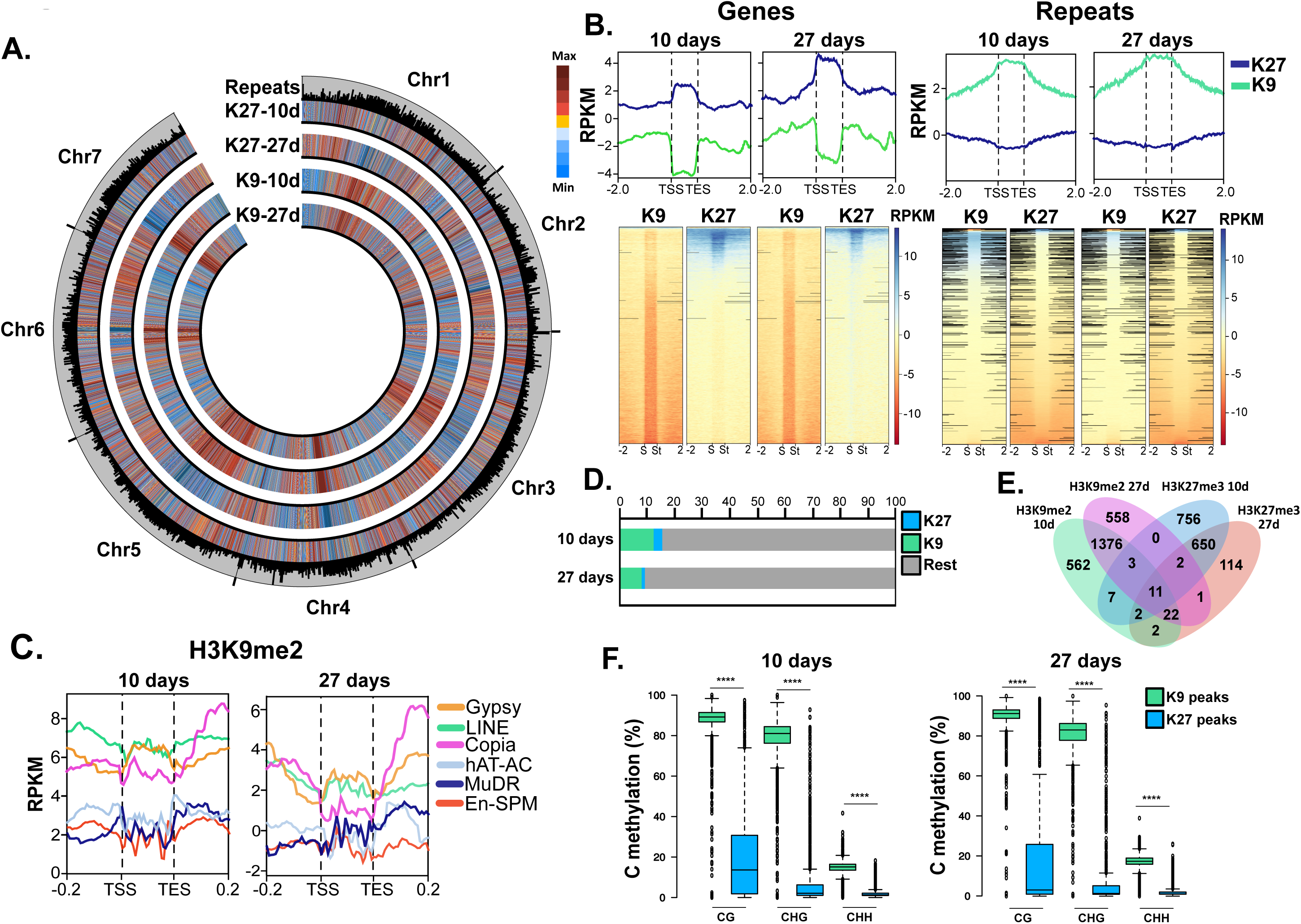
An overview of *Cucumis sativus* heterochromatin. **A.** Circular plot showing the genome-wide levels of H3K27me3 (K27) and H3K9me2 (K9) in mock samples at 10 and 27 dpi. The outermost track indicates the location and length of the repeats present in the *C.sativus* genome. **B.** Histone modification coverage profiles for genes (left) and repeats (right) in mock samples at 10 and 27 dpi. Graphs represent a 2kb window (bottom units) from the transcriptional start site (TSS) and the transcriptional end site (TES). Heatmaps of the values used for the profile graphs are also shown with the transcriptional start site indicated as “S” and the transcriptional end site indicated as “St”. **C.** Histone modification coverage profiles for different TE superfamilies in mock samples at 10 and 27 dpi. Only TEs longer than 2kb were considered for the representation. Graphs represent a 0.2kb window (bottom units) from the transcriptional start site (TSS) and the transcriptional end site (TES). **D.** Percentage of the genome-wide coverage for the peaks identified for each of the histone marks analyzed. **E.** Venn diagram showing the overlapping between the peaks identified in mock tissues for each of the histone marks at each of the time points analyzed. **F.** Values of cytosine methylation percentage for each methylation context for peaks identified on each of the histone marks analyzed. **** indicates a p-value smaller than 0.001. P-values were calculated through an unpaired t-test.

**Figure 2.**
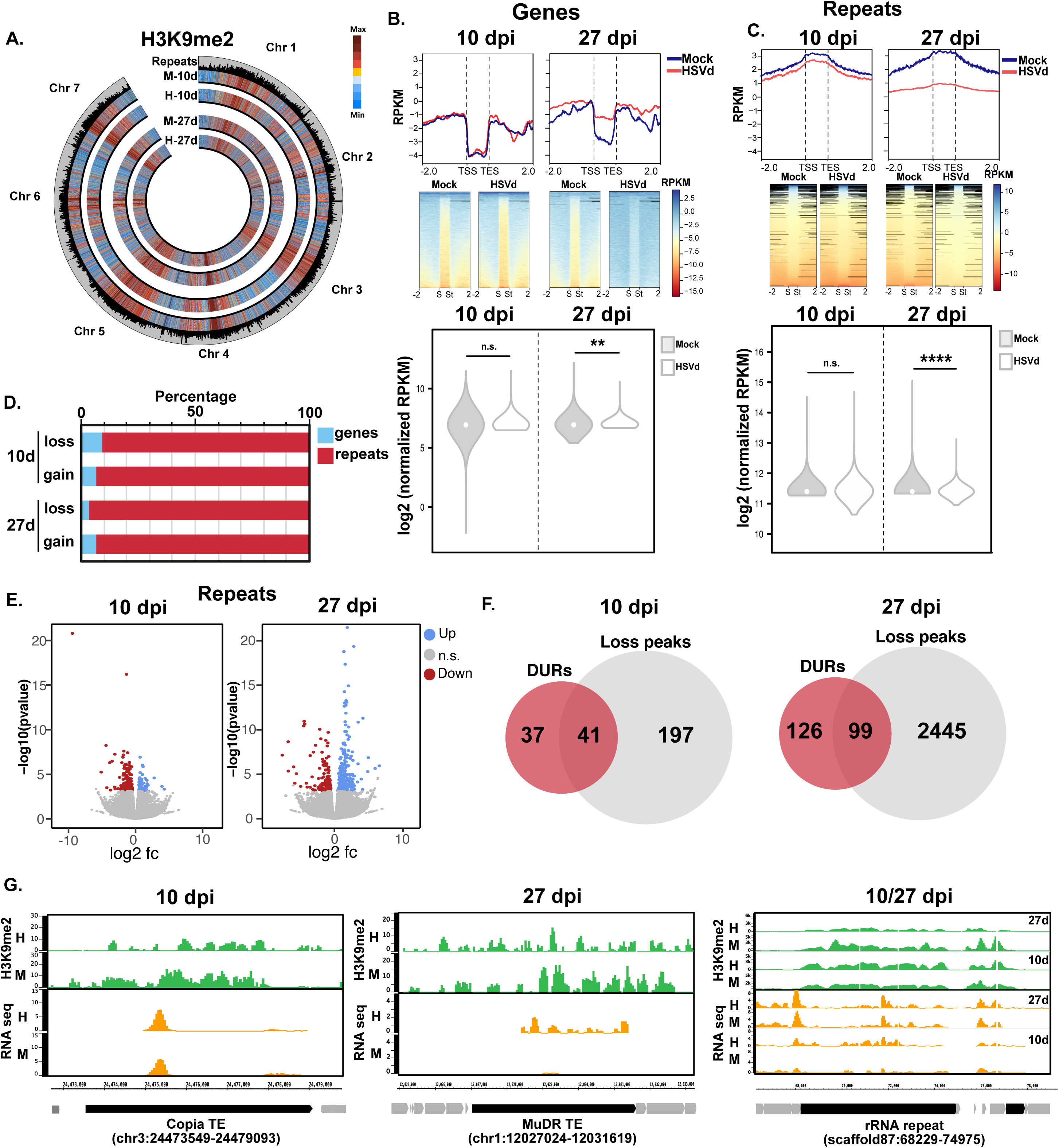
H3K9me2 dynamics under HSVd infection. **A.** Circular plot showing the genome-wide levels of H3K9me2 in mock (M) and HSVD-infected (H) samples at 10 and 27 dpi. The outermost track indicates the location and length of the repeats present in the *C.sativus* genome. **B-C.** Histone modification coverage profiles for genes (**B**) and repeats (**C**) in mock and HSVd-infected samples at 10 and 27 dpi. Graphs represent a 2kb window (bottom units) from the transcriptional start site (TSS) and the transcriptional end site (TES). Heatmaps of the values used for the profile graphs are also shown with the transcriptional start site indicated as “S” and the transcriptional end site indicated as “St”. Profiles are accompanied by a violin plot depicting the distribution of the log 2 transformed normalized RPKM values and the statistical significance of their difference. n.s= no significant, ** and **** indicate a p-value smaller than 0.01 and 0.001 respectively. P-values were calculated through an unpaired t-test. **D.** Percentage of genes (blue) and repeats (red) present in loss or gain peaks identified at each of the time points analyzed. **E.** Volcano plots showing differentially expressed repeats at 10 and 27 dpi. **F.** Venn diagrams showing the overall overlap between 10 and 27 dpi differentially upregulated repeats (DURs) and repeats located at a H3K9me2 loss peak. **G.** Genome browser screenshots showing examples of significantly upregulated repeats located at peaks losing H3K9me2. The time point of the identified overlap is shown on top of the screenshot.

Next, to understand their enrichment at specific locations in the cucumber genome we analyzed the presence of histone peaks (see material and methods for the parameters used for defining peaks) for both marks at the two time points studied. Identification of peaks for the two marks showed that, on average, they occupy a 10% of the cucumber genome (Fig 1D), similar to the inferred percentages from cytogenetic mapping (77). As expected from the different compartmentalization of the marks in genes and repeats (Fig 1B), peaks for the two marks showed little to no overlap (Fig 1E), indicating that they virtually mark two different genomic regions. A characteristic of plant DNA methylation is that overall low DNA methylation levels increase at centromeric and pericentromeric regions (78), where it targets TEs and other repeats (9). Similar to most plant species, there is a significant presence of DNA methylation at H3K9me2 peaks that does not happen at H3K27me3 peaks, suggesting that the RdDM pathway might be directly involved in the establishment of H3K9me2 in cucumber (Fig 1F). In summary, cucumber presented several conserved characteristics in its heterochromatin, including characteristic targeting of TEs by H3K9me2 and genes by H3K27me3 observed in other plant species.

### H3K9me2 and H3K27me3 are reorganized during the progression of HSVd infection

To understand the dynamism experienced by both heterochromatic marks during HSVd infection, we focused on exploring the changes occurring on infected tissues at the two time points under study.

Regarding H3K9me2, at the genome-wide level, HSVd infection did not induce dramatic changes in the distribution of H3K9me2 at any of the two infection points (Fig 2A). Indeed, H3K9me2 changes were only significant at later infection times, where it was gained at genes (Fig 2B) and lost from repeats (Fig 2C). The striking difference in the presence of this mark at repeats compared to genes, indicates again its preferential accumulation at heterochromatic regions and the stability of this enrichment even during HSVd infection (Fig 2B-C). The accumulation changes observed showed that infected plants experienced a re-organization of heterochromatin under HSVd infection at later time points, while this mark was not significantly affected at earlier time points. Next, we predicted H3K9me2 peaks in both mock and infected tissues and explored the gain or loss of peaks of this mark in both conditions and time points (Fig 2D and Sup Table 1). As expected from the preferential heterochromatic nature of this mark, the majority of peaks were located at repeats (94% on average), pointing to the main role of this mark in the control of TEs and other repeats (Fig 2D). To explore this role in regulating repeat transcription, we re-analyzed transcriptomic data obtained at identical time points from a previous work (62). In line with the potential role of H3K9me2 in the transcriptional control of repeats, cucumber plants experience a transcriptional reactivation of repeats at both time points, with a higher number of differentially upregulated repeats (DURs) at later time points (78 and 225 significantly expressed repeats at 10 and 27 dpi respectively, adjusted p-value <0.05, Fig 2E and Sup Table 2). Interestingly, reactivated repeats overlapped with H3K9me2 changes during infection, since 53% and 44% of the reactivated repeats at 10 and 27 dpi, respectively, are within regions that lose H3K9me2 (Fig 2F).

Targets of this reorganization include several TEs from the Copia and MuDR superfamilies (Fig 2G). Importantly, we also observed an important reorganization of H3K9me2 correlated with the transcriptional reactivation of ribosomal RNA repeats (Figure 2G). HSVd is known to induce changes in the DNA methylation levels of ribosomal DNA repeats (46–48,62). Our analysis points to a potential relationship between our previously observed DNA methylation changes at rDNA repeats and the re-organization of H3K9me2 during HSVd infection, since both marks are intrinsically connected in cucumber (Fig 1F). Despite this, the majority of H3K9me2 changes were not connected with DNA methylation changes, since almost all peaks retained similar DNA methylation values between mock and infected samples (the exception being CHH values in H3K9me2 gain peaks at 10 dpi which significantly decreased, Sup Fig 3A).

**Figure 3.**
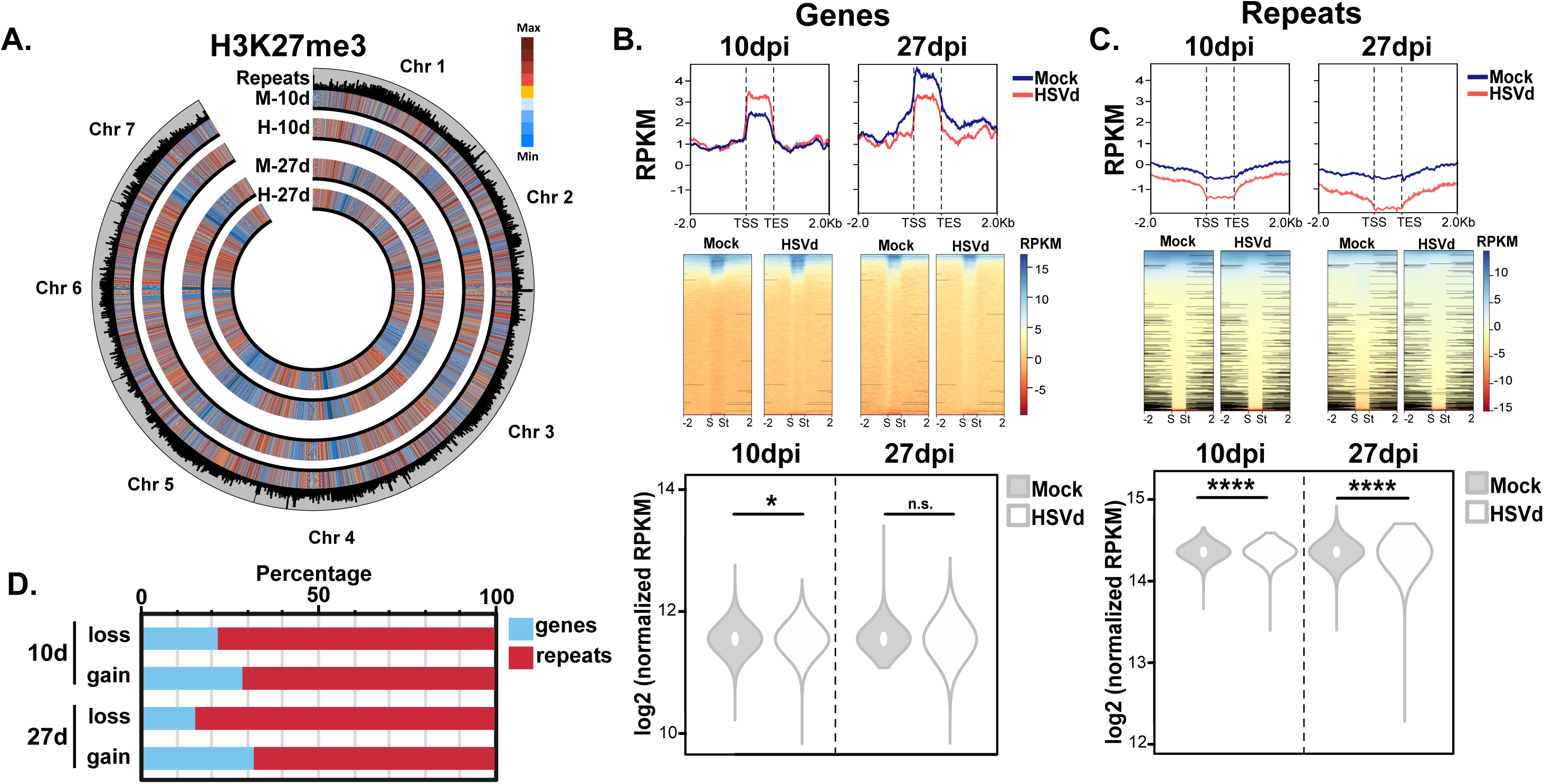
H3K27me3 dynamics under HSVd infection. **A.** Circular plot showing the genome-wide levels of H3K27me3 in mock (M) and HSVD-infected (H) samples at 10 and 27 dpi. Outermost track indicates the location and length of the repeats present in the *C.sativus* genome. **B-C.** Histone modification coverage profiles for genes (**B**) and repeats (**C**) in mock and HSVd-infected samples at 10 and 27 dpi. Graphs represent a 2kb window (bottom units) from the transcriptional start site (TSS) and the transcriptional end site (TES). Heatmaps of the values used for the profile graphs are also shown with the transcriptional start site indicated as “S” and the transcriptional end site indicated as “St”. Profiles are accompanied by a violin plot depicting the distribution of the log 2 transformed normalized RPKM values and the statistical significance of their difference. n.s= no significant, * and **** indicate a p-value smaller than 0.05 and 0.001 respectively. P-values were calculated through an unpaired t-test. **D.** Percentage of genes (blue) and repeats (red) present in loss or gain peaks identified at each of the time points analyzed.

Interestingly, H3K27me3 also followed a discrete re-organization under HSVd infection (Fig 3A). At the genome-wide level, this mark showed a significant enrichment in genes only at the earlier infection time point (Fig 3B), while it was significantly lost from repeats at both 10 and 27 dpi (Fig 3C). Similar to our H3K9me2 analysis, we predicted H3K27me3 peaks in both mock and infected tissues and explored the gain or loss of peaks of this mark in both conditions and time points (Fig 3D and Sup Table 3). Compared to H3K9me2 (6% on average, Fig 2D), H3K27me3 peaks are more frequents at genic locations (24% on average, Fig 3D). Despite the presence of some of the peaks at repeat locations, their overall values (Fig 3C) indicate that the presence of this mark at repeats is anecdotal. As expected, changes observed for this mark are independent of DNA methylation (Sup Fig 3B). Overall, HSVd infection induced a reorganization of constitutive and facultative heterochromatin that was connected to the transcriptional reactivation of repeats, including TEs and rRNA genes.

### H3K9me2 and H3K27me3 retain constitutive and facultative heterochromatic identity during viroid infection

We focused on understanding the dynamics of both heterochromatic marks under HSVd infection. First, we analyzed the dynamism of our predicted gain and loss peaks (Fig 4A). Overall, on average, 20% of H3K9me2 and 31% of H3K27me3 peaks showed a dynamic nature (gain or loss of the peak under HSVd infection) while the rest (80 and 69% of the H3K9me2 and H3K27me3 peaks, respectively) remained stable. Following their main occupancy preference, gain and loss of the marks took place in centromeric/pericentromeric regions (for H3K9me2) and the arms of the chromosomes for H3K27me3 (Fig 4B).

**Figure 4.**
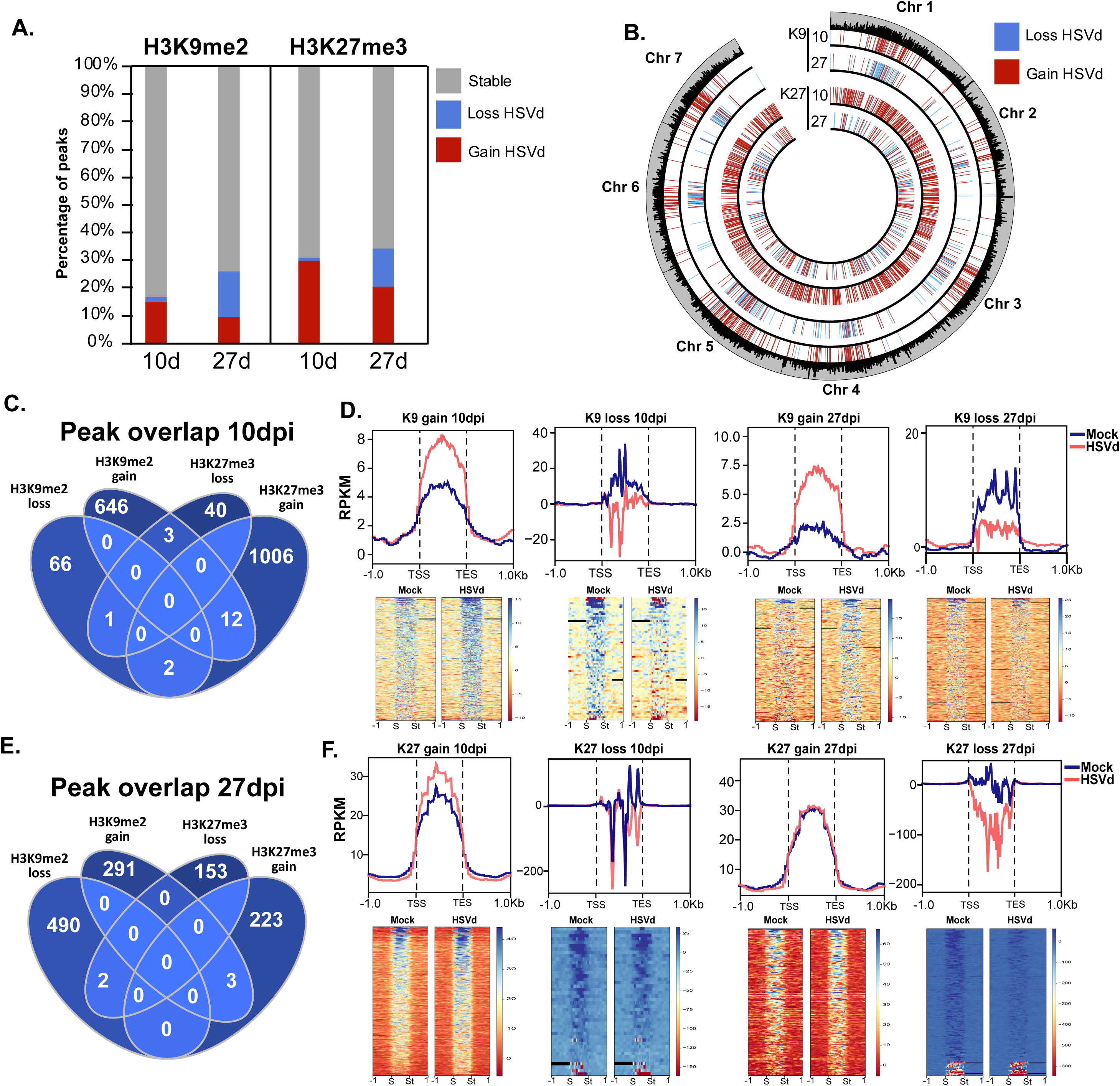
Genome-wide analysis of H3K9me2 and H3K27me3 changes under HSVd infection. **A.** Analysis of the dynamism of the peaks identified under HSVd infection for both H3K9me2 and H3K27me3. Stable peaks were identified in both mock and HSVd-infected samples, while loss and gain peaks were only identified in mock or HSVd-infected samples, respectively. **B.** Circular plot showing the genome-wide location of the identified loss and gain histone peaks for both H3K9me2 (K9) and H3K27me3 (K27) samples at both 10 and 27 dpi. Outermost track indicates the location and length of the repeats present in the *C.sativus* genome. **C.** Venn diagram showing the overlap between the H3K9me2 and H3K27me3 gain and loss peaks at 10 dpi. **D.** Histone modification coverage profiles for the identified H3K9me2 peaks in mock and HSVd-infected samples at 10 and 27 dpi. Graphs represent a 2kb window (bottom units) from the transcriptional start site (TSS) and the transcriptional end site (TES). Heatmaps of the values used for the profile graphs are also shown with the transcriptional start site indicated as “S” and the transcriptional end site indicated as “St”. **E.** Venn diagram showing the overlap between the H3K9me2 and H3K27me3 gain and loss peaks at 27 dpi. **F.** Histone modification coverage profiles for the identified H3K27me3 peaks in mock and HSVd-infected samples at 10 and 27 dpi. Graphs represent a 2kb window (bottom units) from the transcriptional start site (TSS) and the transcriptional end site (TES). Heatmaps of the values used for the profile graphs are also shown with the transcriptional start site indicated as “S” and the transcriptional end site indicated as “St”.

H3K9me2 and H3K27me3 are the main histone marks delimiting plant heterochromatin (79). In Arabidopsis, H3K27me3 can “invade” constitutive heterochromatic regions under heavy loss of DNA methylation and/or heterochromatic identity in mutants, or under viral stress (37,80,81). Interestingly, despite the overall reprogramming of the two marks observed during viroid infection (Fig 2 and Fig 3), there was a minimal interaction (if any) between the loss and gain of any of the repressive histone marks (Fig 4C and Fig 4E). Analysis of the accumulation values for both marks showed interesting trends. First, at both analyzed times, H3K9me2 was gained in regions with average/low mark values, while it was lost at locations showing a higher enrichment (Fig 4D). This result correlated with our observed effect of H3K9me2 loss over the transcriptional reactivation of repeats (which were located at H3K9me2 rich regions). On the other hand, H3K27me3 was gained (although at lower levels) at regions with high values of the mark, while it was lost at regions with low values (Fig 4F). Therefore, re-organization of heterochromatin under HSVd infection was limited to regions with heterochromatic identity.

### Changes in gene expression correlate with H3K27me3 gain

Next, we aimed to understand the influence of H3K9me2 and H3K27me3 re-organization on the genic transcriptional response to HSVd infection. First, we analyzed the overlap of genes with peaks showing a dynamic behavior under infection. We restricted our analysis to: 1) genes directly associated with the gain or loss of H3K27me3 within their gene bodies, and 2) genes that gained or lost H3K9me2 within their gene bodies and a ± 1kb window from their gene bodies.

Using this strategy, we identified 310 and 1,221 genes with expression associated to changes in H3K9me2 and H3K27me3 marks, respectively (Fig 5A and Sup Table 4). Most of the genes (67% and 79%, for H3K9me2 and H3K27me3, respectively) were associated with gain peaks. To understand the role of genes regulated by repressive histone marks we explored their gene ontology (GO) classification. Genes overlapping with these marks were mainly associated with DNA, protein and RNA binding categories, and with the transferase, hydrolase, and catalytic activities categories (Figure 5B). Interestingly, some of these categories were preferentially associated with each of the marks. For example, DNA binding and kinase activity were predominantly associated with H3K27me3, while hydrolase and catalytic activities were in general related to H3K9me2 (Figure 5B). This observation is in line with the known roles of H3K27me3 in regulating the sensing of environmental signals (82) and a potential role of H3K9me2 in the control of housekeeping metabolic activities as observed in other species such as *C.elegans* or humans (83,84). Next, we explored the influence of the presence of gain or loss peaks over the transcriptional activation or repression of gene expression in response to HSVd infection. To that end, we used transcriptional data from previous experiments at analogous time points to infer the expression level of the genes located within histone peaks. Our analysis indicated that from the different marks and their respective environments, only the gain of H3K27me3 significantly overlapped with the overall decrease of the expression of its target genes at both analyzed time points (Fig 5C).

**Figure 5.**
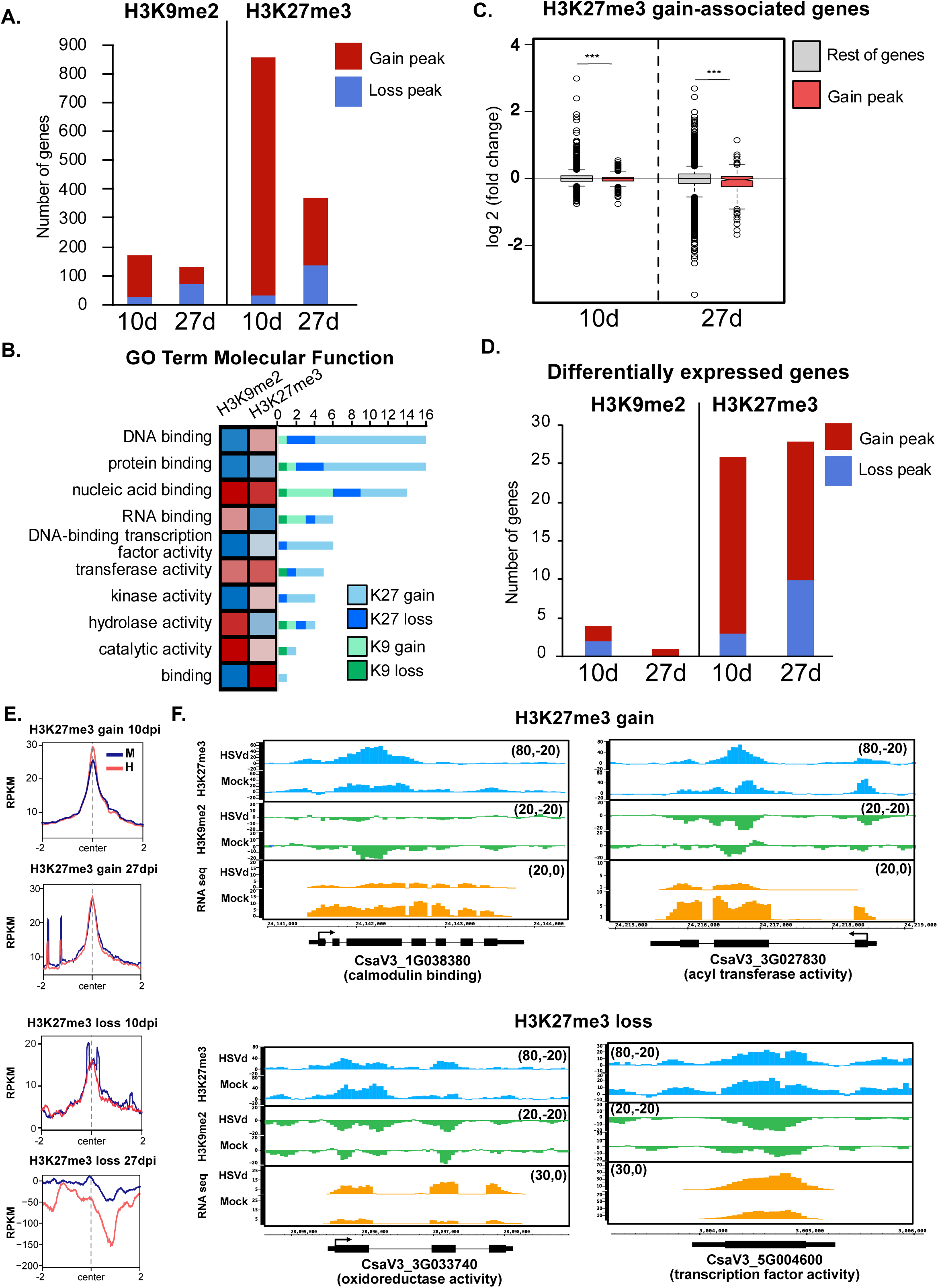
H3K27me3 and H3K9me2 contribute to the transcriptional reprogramming under HSVd infection. **A.** Histogram showing the total number of genes located at gain (red) and loss (blue) peaks for H3K9me2 and H3K27me3 at 10 and 27 dpi. **B.** Gene ontology (GO) categorization by molecular function for the categories indicated of the genes showing association with either H3K9me2 or H3K27me3 peaks. Heatmap shows the enrichment of the terms for peaks identified associated with either H3K9me2 or H3K27me3 peaks. Histogram showing the number of genes associated with gain or loss peaks for each of the histone marks analyzed. **C.** Expression values (log 2 (fold change)) for all genes associated with H3K27me3 gain peaks at 10 and 27 dpi (colored in red) compared to the rest of the genes (all genes with expression in the *C.sativus* genome with the subtracted values for peak-associated genes). p-values are indicated on top of each comparison and were calculated through an unpaired t-test. ***= p-value lower than 0.005. **D.** Histogram showing the total number of differentially expressed genes located at gain (red) and loss (blue) peaks for H3K9me2 and H3K27me3 at 10 and 27 dpi. **E.** H3K27me3 coverage profiles for the indicated H3K27me3 peaks in mock (M) and HSVd-infected (H) samples at 10 and 27 dpi. Graphs represent a 2kb window (bottom units) from the center of the region. **F.** Genome browser screenshots showing examples of significantly upregulated repeats located at peaks gaining or losing H3K27me3.

Several differentially expressed genes (DEGs, 26 at 10dpi and 28 at 27dpi) were found associated with H3K27me3 changes, with both gain (23 at 10dpi and 18 and 27dpi) and loss (3 at 10dpi and 10 at 27dpi) of marks (Fig 5D and Sup Table 4). Interestingly, the gain of H3K27me3 took place at genes with already high values of H3K27me3. In contrast, loss of this mark took place at genes with medium/low values of the mark, indicating that this mark is not gained *de novo* but rather, reinforced at previous heterochromatic locations (Fig 5E). DEGs associated with H3K27me3 peaks included genes potentially involved in the mediation of the stress-response. For example, expression of calmodulin binding (CsaV3_1G038380) and acyl-transferase activity (CsaV3_3G027830) genes (associated with H3K27me3 gain) or an oxidoreductase (CsaV3_3G033740) and transcription factor (CsaV3_5G004600) genes (Fig 5F). Additionally, certain DEGs (4 at 10dpi and 1 at 27dpi) were also associated with H3K9me2 changes, including CsaV3_ 7G025330, a bystin-like protein, potentially involved in cell proliferation (85,86), and CsaV3_7G015110, a homolog of the enhanced downy mildew 2 protein, a NLS protein that may regulate RPP7 expression (87), which has previously been proposed to be epigenetically regulated by heterochromatin in *Arabidopsis* (88) (Sup Fig 4). Overall, our data indicate a role of histone marks in the regulation of the transcriptional response to HSVd infection with a particular main role of the repressive character of H3K27me3 and a minor contribution of H3K9me2 in the control of gene expression.

## DISCUSSION

Epigenetic regulation plays an important role in the adaptation of the transcriptional program to external signals that determine plant development (22–25,28). Stress associated to adverse environmental conditions has been identified as yet another stimulus that can induce epigenetic changes in plants (6,7,89). Several studies have identified dynamic DNA methylation changes upon stress exposure, but the role of histone marks in this context has remained unexplored. In this work, we have studied the dynamism and role of two repressive histone marks (H3K9me2 and H3K27me3) in the modulation of the transcriptional response against viroid infection. Our data shows that heterochromatic histone marks are reorganized during HSVd infection, with a major role of H3K27me3 gain in the repression of gene expression.

Our work adds yet another piece of evidence to the role of histone marks in the control of gene expression in response to stress conditions, which have been identified as major players in forward genetic analysis, but were missing genome-wide information of this role. Additionally, our analysis contributes to the understanding of the epigenomic organization of economically relevant crops, with our analysis of the presence of repressive histone marks in the *Cucumis sativus* genome.

The organization of repressive histone marks in the cucumber genome resembles their previously described distribution in *Arabidopsis* (76), melon (90,91), and *Physcomitrella patens* (92), with H3K9me2 marking constitutive heterochromatin (located at the chromosomal centromeric regions) and H3K27me3 featuring facultative heterochromatin (spaced through the chromosome arms). Our results also correlate with previous cytogenetic analyses of cucumber chromatin (77). In line with this preferential accumulation, H3K27me3 and H3K9me2 are highly enriched at genes and repeats, respectively. Similar to *Arabidopsis*, H3K9me2 in cucumber is also correlated with the presence of DNA methylation, since H3K9me2 enriched regions showed high values of cytosine methylation in all sequence contexts. This points to a connection between the RdDM pathway and H3K9me2 homeostasis in cucumber, similar to the one inferred from the genetic analysis of CMT3 and KYP mutants in tomato (93) and pointing to a conservation of that connection in angiosperms.

HSVd-infection induces a global re-organization of cucumber heterochromatin characterized by a progressive reduction of H3K9me2 from repeats and an initial increase followed by a decrease of H3K27me3 from genes. Changes in the presence of both histone marks are influenced by the previous presence of those marks at the enriched peaks, with most of the gain and loss of H3K9me2 and H3K27me3 gain taking place at regions already enriched on that particular mark. On the other hand, H3K27me3 loss seems to take place only in regions with low values of that mark. In line with this accumulation pattern and the canonical role of both repressive marks, repeats are transcriptionally reactivated at later infection points, while genes are mostly affected by gain of H3K27me3. Similar results were recently obtained in virus-infected plants, where a catalytic component of the PRC2 complex (CURLY LEAF) was identified as a factor promoting viral tolerance (37), and in general H3K27me3 homeostasis is an important mark regulating both the stress response (38,92) and developmental cues (22–28).

Previous analysis of the interaction between the repressive histone marks H3K9me2 and H3K27me3 have identified a compensatory mechanism where H3K27me3 invades H3K9me2 regions in mutant backgrounds that lose constitutive heterochromatin (80,81) or under CMV infection (37). Here we were not able to identify a similar mechanism, an indeed, despite the strong symptomatology displayed by plants infected with HSVd (similar to CMV) the two heterochromatic marks retained their identity. Further analysis under different/stronger stresses would be needed to assess if H3K9me2/H3K27me3 compensatory activity is exclusive of the *Arabidopsis* genome or is also present in other plant genomes.

Lastly, we correlated the observed changes in repressive histone marks to the transcriptional changes previously associated with HSVd infection (62). This analysis indicated that, as expected (76), H3K27me3 is vastly more associated with genes than H3K9me2. Nevertheless, we identified some genes whose expression correlate with the presence of enriched or depleted regions for each of the histone mark. Additionally, some of the gene categories seem to be preferentially associated with each of the two repressive histone marks. Further analysis on the connection between histone marks and particular gene families/categories should be carried out to confirm this observation. Furthermore, compared to H3K9me2, H3K27me3 is preferentially associated with DEGs, supporting the role of this mark in the transcriptional reprogramming under HSVd infection. Indeed we identified several genes associated with the gain or loss of this mark that might be important regulators of HSVd defense response, including a calmodulin binding gene, which mediate the first wave of plant responses to multiple stresses through Ca^2+^ binding (94–96), an acyl-transferase gene, which are known to affect plant growth (97,98), an oxidoreductase activity gene, known for controlling oxidative stress in plants (99), or a transcription factor, among other DEGs.

How could a minimal pathogen such as a viroid induce these dramatic changes in histone homeostasis? We envision that this interaction might be indirect, due to the transcriptomic reprogramming needed to cope with the infection, or direct, through interaction with components of histone homeostasis (Sup Fig 5). HSVd is a member of the *Pospiviroidae* family, characterized by their ability to replicate in the nucleus. It is plausible that the accumulation of HSVd in the nucleus alters the homeostasis of that organelle environment, inducing changes in the epigenetic regulation of the genome at its core, the structure of the chromatin. Indeed, HSVd interacts with HDA6, a histone deacetylase that is needed to enhance DNA methylation mediated by MET1 and histone modifications mediated by the histone demethylase FLD (an LSD1 homolog) (100,101). FLD functions as an activator of flowering through the repression of FLC, which is controlled by H3K27me3 (102). This interaction with histone homeostasis could lead to the increased values of H3K27me3, which seems of particular importance under HSVd infection-induced transcriptional reprogramming. Indeed, HSVd-infected plants show a delay in flowering time (103), which could be connected to our observed H3K27me3 reorganization. Since in our analyses, the changes observed at H3K9me2 peaks were independent of any DNA methylation change, we disfavor that HSVd interactions with histone modifications is a consequence of the interference of HSVd-derived siRNAs with the RdDM pathway and, consequently, H3K9me2 homeostasis. To summarize, our work adds data supporting the role of epigenomic reprogramming, and particularly heterochromatin reorganization, playing an active role in the transcriptional reprogramming experienced under viroid infection. Moreover, our results expand our understanding of the complex interaction between viroids and their hosts.

## AUTHOR CONTRIBUTIONS

Joan Marquez-Molins: Formal analysis, Methodology, Validation, Writing—review & editing. Jinping Cheng: Formal analysis, Methodology, Validation, Writing—review & editing. Julia Corell-Sierra: Bioinfomatic Analysis. Vasti Thamara Juarez-Gonzalez: Formal analysis, Methodology, Bioinfomatic Analysis, Writing—review & editing. Pascual Villalba-Bermell: Bioinfomatic Analysis. Maria Luz Annacondia: Formal analysis, Methodology, Validation, Writing—review & editing. Gustavo Gomez: Conceptualization, Formal analysis, Writing— review & editing. German Martinez: Conceptualization, Formal analysis, Methodology, Validation, Bioinfomatic Analysis, Writing—original draft.

## ACKNOWLEDGEMENTS

We thank all members of the Martinez and Gomez labs for valuable comments on the manuscript. Sequencing was performed by Novogene (United Kingdom). The data handling was performed using UPPMAX which is part of the Swedish National Infrastructure for Computing (SNIC).

## FUNDING

We thank Formas (2021–01161), the Swedish Research Council (VR 2021-05023), and the Knut and Alice Wallenberg Foundation (KAW 2019.0062) for supporting research in the Martinez group, and The Knowledge Generation Program of the Spanish Research Agency (PID2022_1393930B-I00) in the Gomez group. Open Access funding provided by Swedish University of Agricultural Sciences. The data handling was enabled by resources provided by the Swedish National Infrastructure for Computing (SNIC) at UPPMAX partially funded by the Swedish Research Council through grant agreement no. 2018-05973.

## CONFLICT OF INTEREST

None declare

## Notes

### Competing Interest Statement

The authors have declared no competing interest.

