## Supplementary figures and images for "Hop stunt viroid infection alters host heterochromatin"

### Supplementary Figure 1

#### Supplementary Figure 1

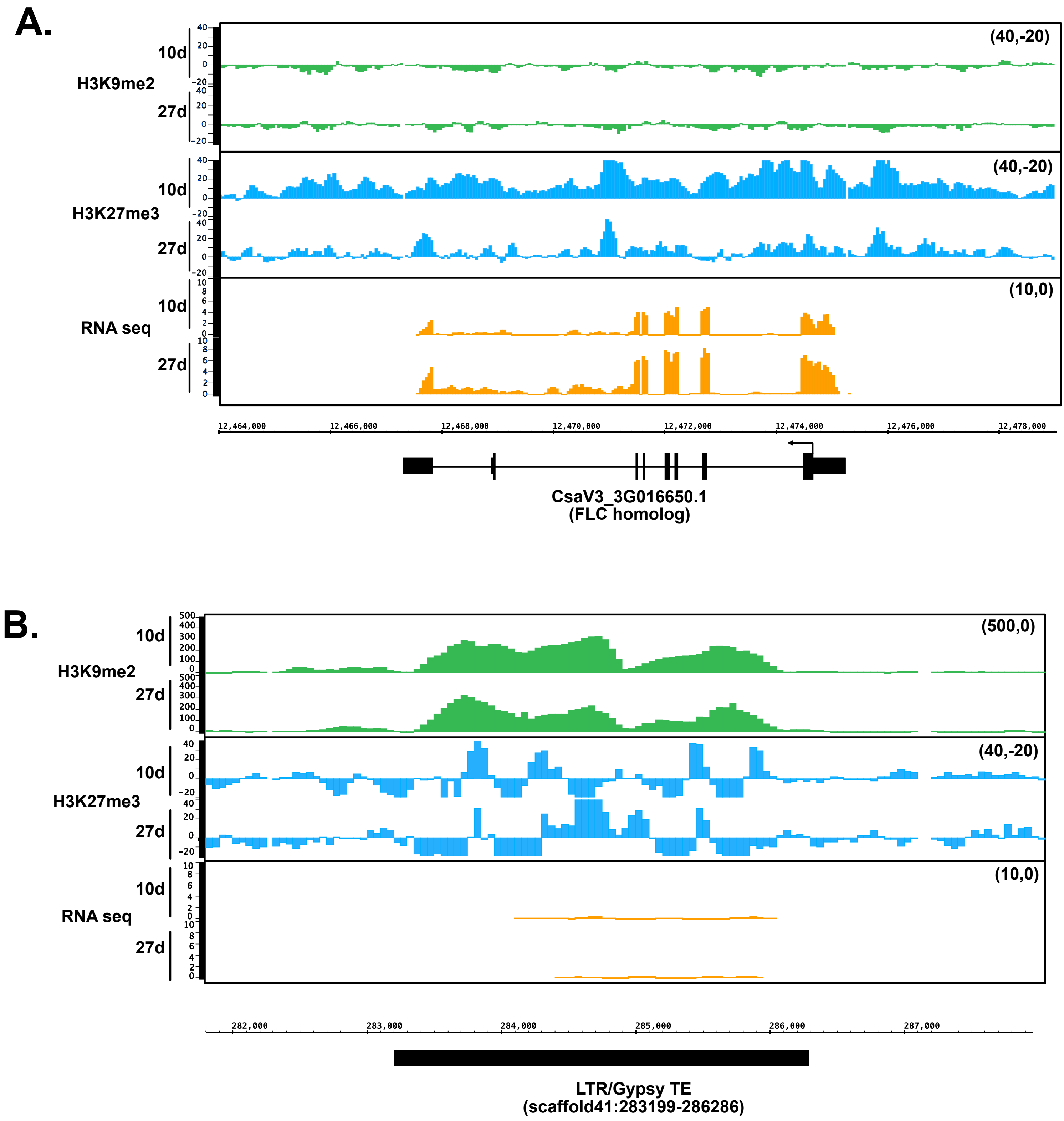

### Supplementary Figure 2

#### Supplementary Figure 2

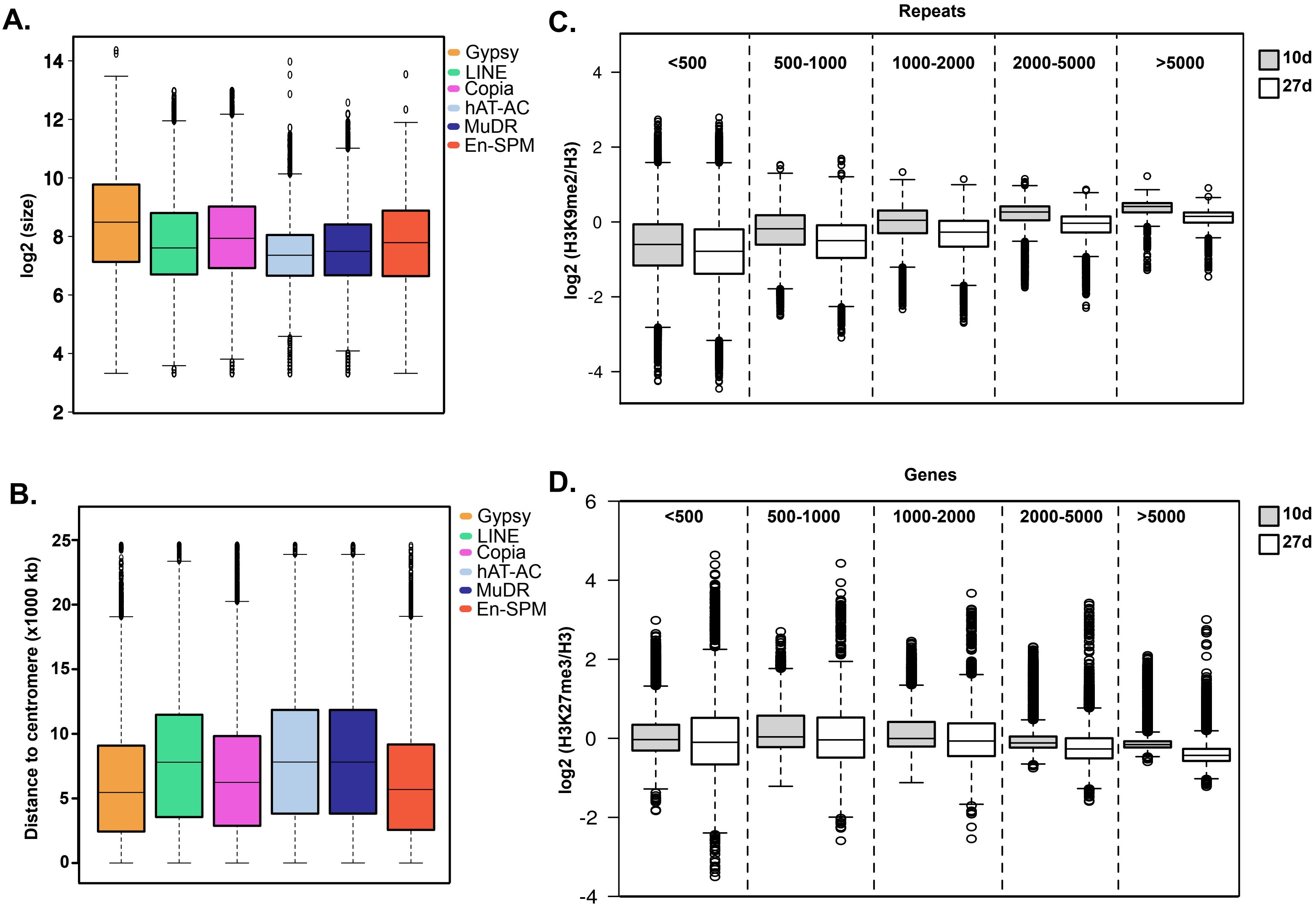

### Supplementary Figure 3

#### Supplementary figure 3

A.

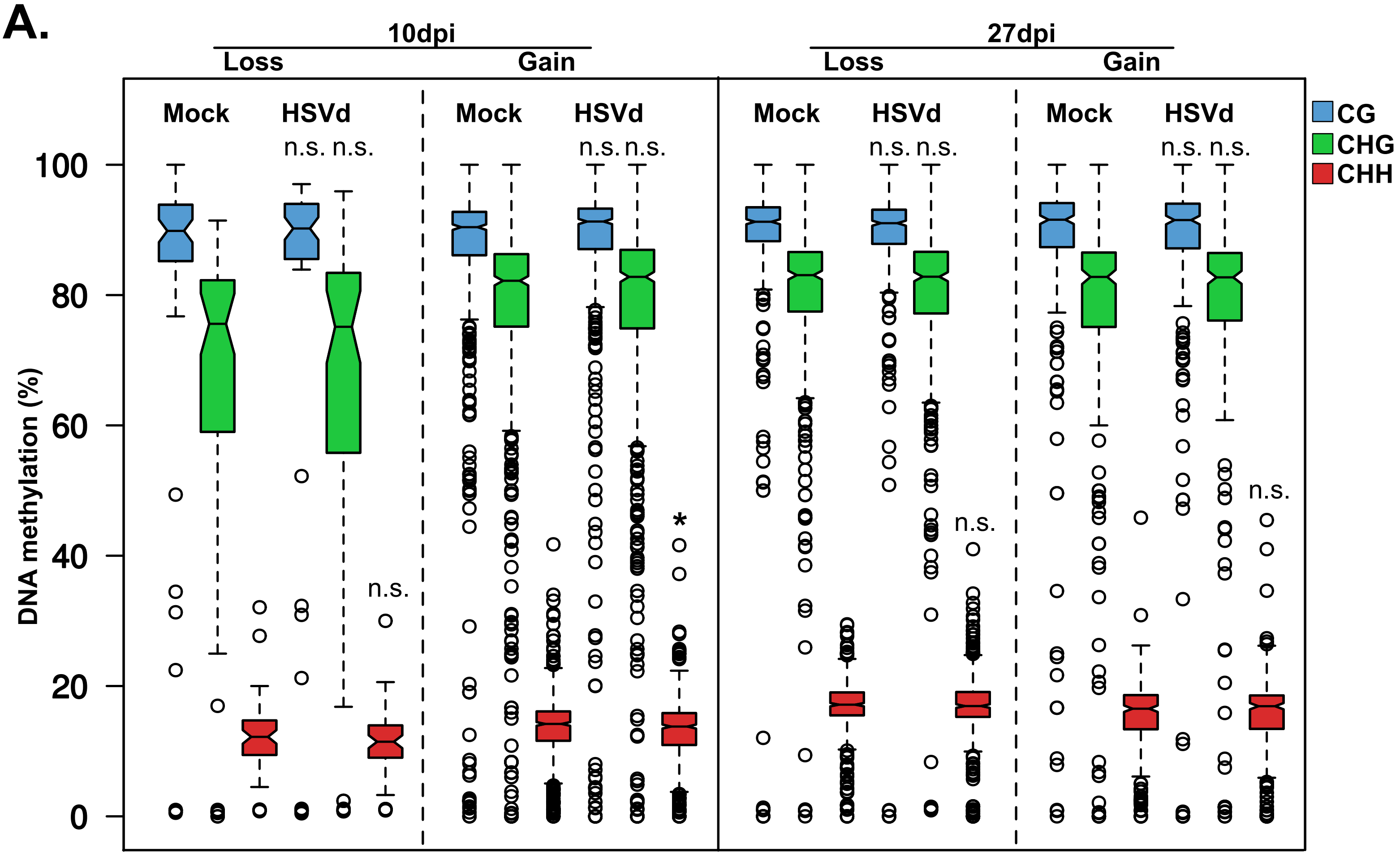


B.

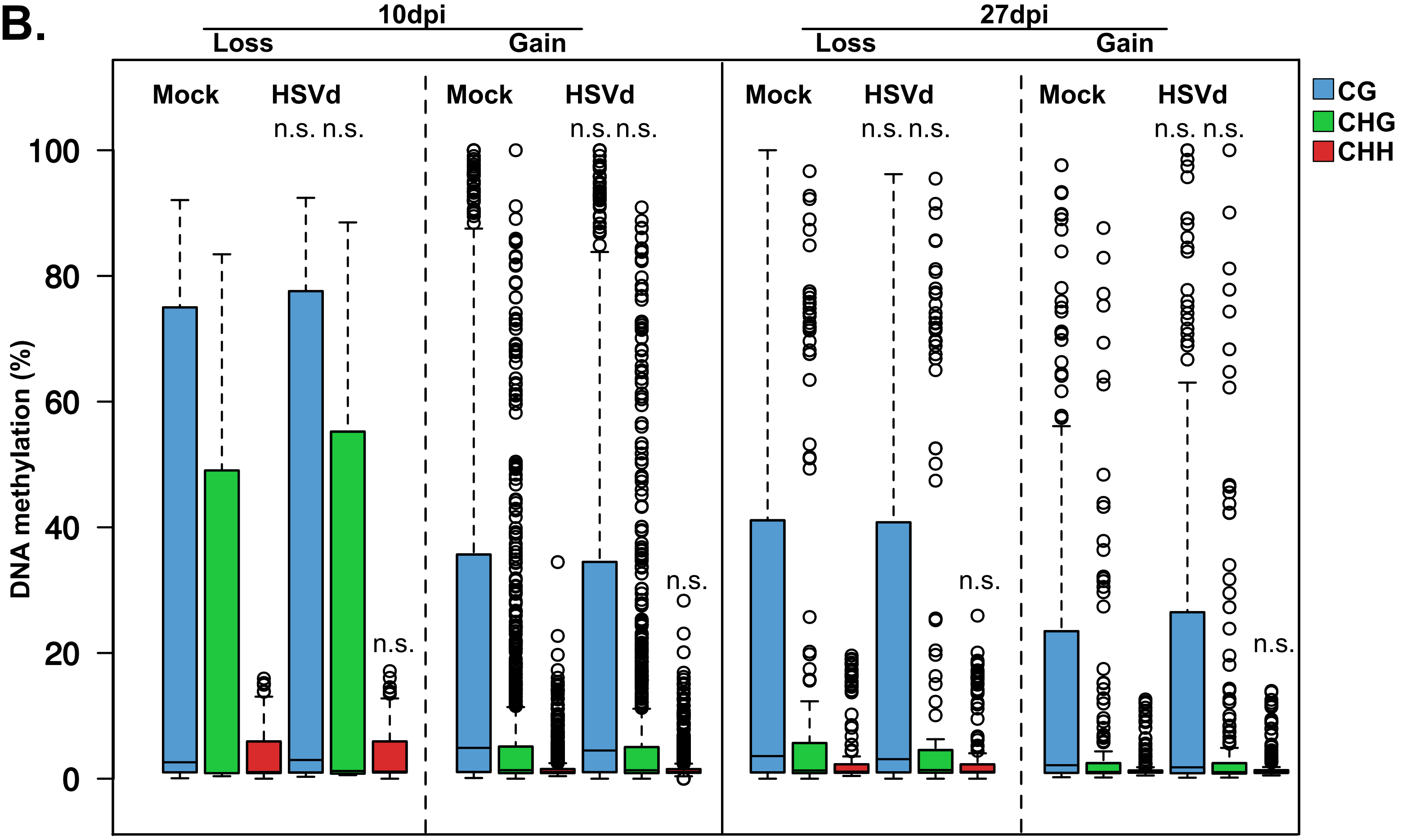

### Supplementary Figure 4

#### Supplementary figure 4

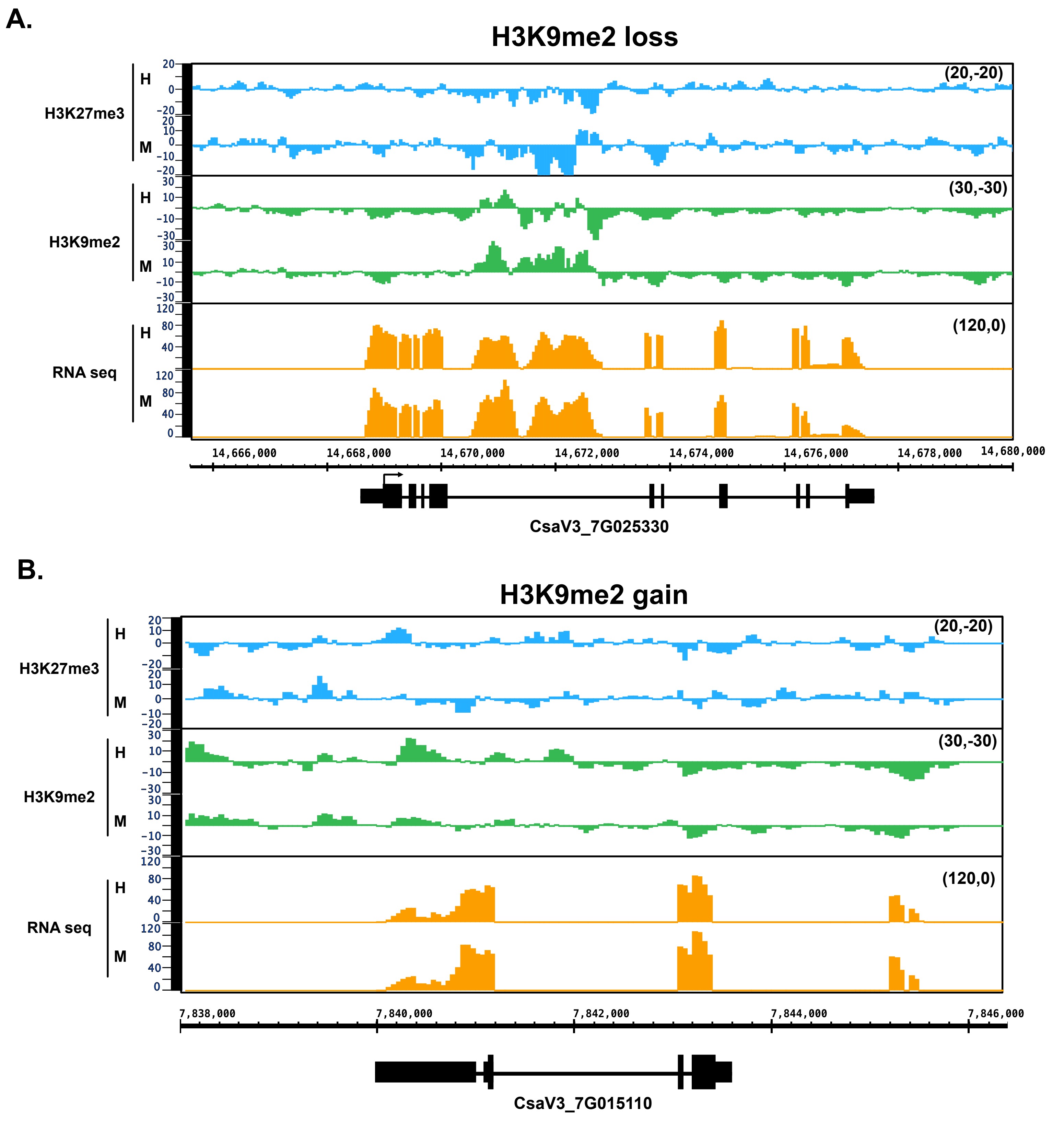

### Supplementary Figure 5

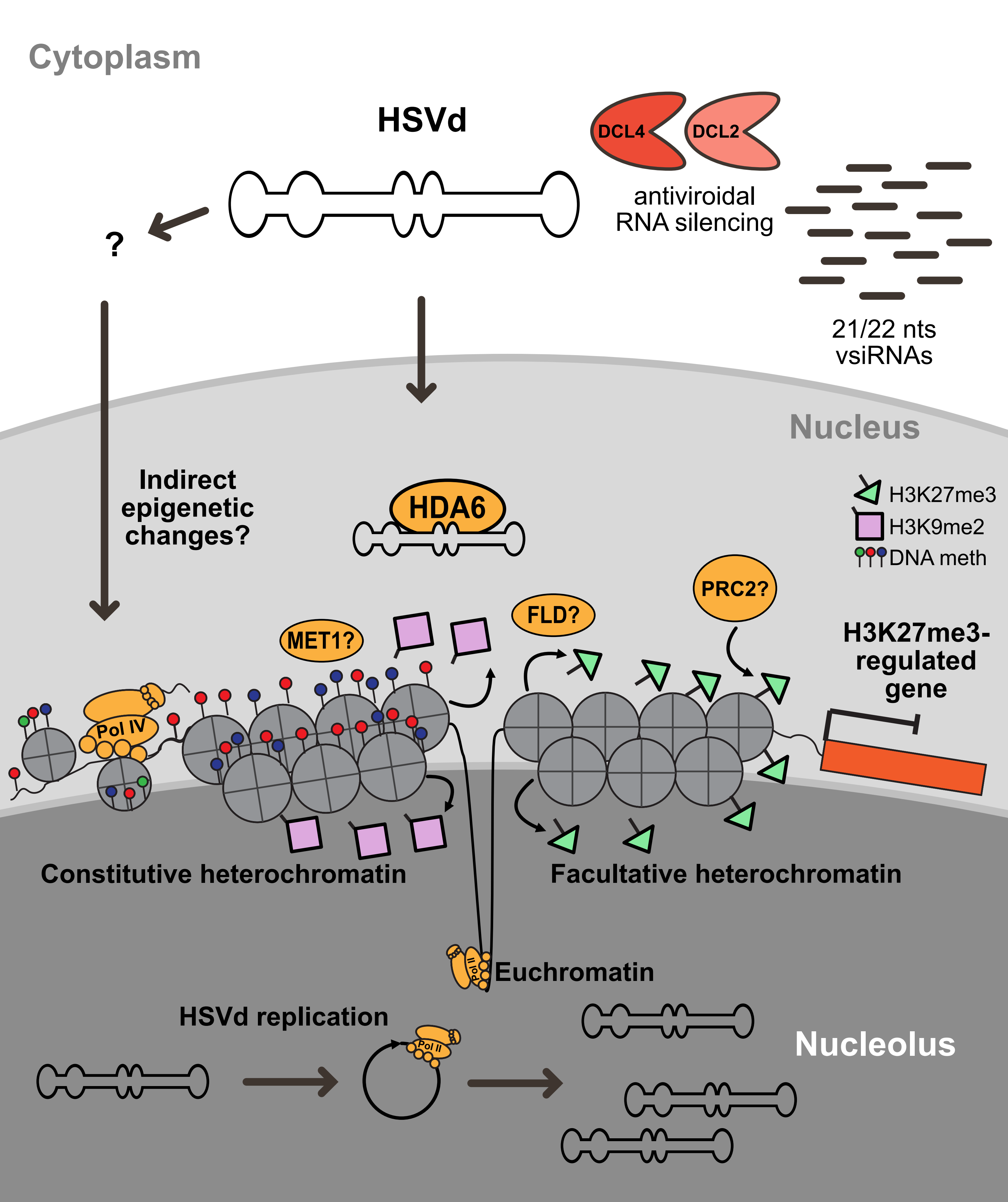
